# Drospondin, a novel glial-secreted glycoprotein, controls the development and function of the *Drosophila* nervous system

**DOI:** 10.1101/2024.11.08.622737

**Authors:** Francisca Rojo-Cortés, Candy B. Roa-Siegfried, Nicolás Fuenzalida-Uribe, Paula Amado-Hinojosa, María-Constanza González-Ramírez, Isidora Almonacid-Torres, Serge Birman, Lindsey D. Goodman, Oguz Kanca, Carlos Oliva, María Paz Marzolo, Jorge M. Campusano

## Abstract

Reelin is a secreted glycoprotein with roles in the development of the mammalian neocortex, hippocampus and cerebellum. This vertebrate signaling molecule also contributes to adult brain function. Mammalian Reelin increases the complexity of *Drosophila* Mushroom Body (MB) neurites, an effect mediated by LpR1 and LpR2, the orthologs of mammalian Reelin receptors. Paradoxically, to date, no Reelin ortholog has been described in *Drosophila*. Here, we report that the protein product of the uncharacterized *Drosophila CG17739* gene, which we named Drospondin, shares sequence homology with vertebrate F-spondin and Reelin. We show that Drospondin is expressed in glial cells and is crucial for MB development. Our results also show that Drospondin genetically interacts with LpRs and that human Reelin rescues structural defects in Drospondin-deficient flies. Furthermore, Drospondin-deficient flies exhibit altered sleep homeostasis, locomotion, and social behaviors. Our results reveal that flies express a functional homolog of mammalian Reelin that controls the development and function of the *Drosophila* nervous system.

## INTRODUCTION

Diverse signaling molecules play critical roles in several stages of neuronal development. Importantly, mutations in these molecules or the cellular components associated with their signaling contribute to anatomical or functional alterations that might underlie neurodevelopmental or neurodegenerative disorders.

Reelin is an extracellular glycoprotein with crucial functions in the vertebrate central nervous system (CNS), controlling the radial migration and differentiation of neurons in laminated structures, including the hippocampus, cerebellum and cerebral cortex [1–3], as well as the migration of mesencephalic dopaminergic neurons [4]. In adulthood, Reelin has additional roles in synaptic communication and plasticity by modulating α-amino-3-hydroxy-5-methyl-4-isoxazolepropionic acid (AMPA) and N-methyl-D-aspartate (NMDA) receptor activity, upon activation of its receptors apolipoprotein E receptor 2 (ApoER2) and very-low-density-lipoprotein receptor (VLDLR) [5]. Reelin also contributes to synaptic communication by the enhancement of spontaneous neurotransmitter release [6]. Mice deficient in Reelin, ApoER2, or VLDLR exhibit defects in long-term potentiation and memory formation [7, 8]. Mutations in the Reelin gene, *RELN,* have been associated with lissencephaly [9]. GWAS and linkage studies support that they are also linked to some psychotic and mood disorders, autism spectrum disorder (ASD), schizophrenia, and Alzheimer’s disease (AD) [10–14]. Reelin is expressed by Cajal-Retzius cells located in the Marginal Zone in the developing CNS [3] and in GABAergic interneurons in the adult brain [3]. Reelin is also secreted by Schwann cells and activates their migration, thus contributing to axonal regeneration and outgrowth at the periphery [15, 16].

We previously demonstrated that cultured *Drosophila* Mushroom Body (MB) neurons respond to mammalian Reelin by increasing the complexity of their neurites [17], similar to what is observed for neurons from the mammalian hippocampus and cerebral cortex [18]. We also showed that Reelin-induced responses in cultured *Drosophila* neurons depend on Lipophorin receptor 1 (LpR1) and Lipophorin receptor 2 (LpR2) [17]. *Drosophila* LpR1 and LpR2 are two proteins with high similarity in amino acid sequence and domain organization to VLDLR and ApoER2 [19, 20]. Furthermore, Reelin-induced effects on *Drosophila* neurons depend on disabled (Dab) [17], the fly ortholog of Dab1, which is the intracellular adaptor activated by ApoER2 and VLDLR in the Reelin signaling cascade in vertebrates [3].

The MB is a crucial structure in the fly brain, which has been implicated in olfactory learning and memory, locomotor activity and sleep regulation, among other functions [21–23]. Recently, we demonstrated phenotypes in sleep dynamics, and olfactory learning and memory in LpR-deficient flies [17]. Furthermore, we also showed that LpRs participate in axonal guidance in the MB. Thus, our data support that the axon pathfinding defects detected in the MB of LpR-deficient flies have consequences for adult MB structure and function [17]. To date, no Reelin-like protein has been described in *Drosophila*. Moreover, it has been historically thought that Reelin signaling is only present in vertebrates [24]. Interestingly, a genetic analysis concluded that proteins that could have similar functions to vertebrate Reelin appeared in evolution with the phylum Arthropoda [25], a group of animals to which *Drosophila melanogaster* belongs. Given that *Drosophila* has orthologs for proteins involved in the vertebrate Reelin signaling pathway, including the receptors VLDLR and ApoER2, we hypothesized that a Reelin-like pathway must exist in *Drosophila* [17].

Here we tested the hypothesis that a protein with similar functions to Reelin is encoded in the *Drosophila* genome. We identified a previously uncharacterized protein encoded by the *CG17739* gene, with the highest amino acid similarity to Reelin. Because the predicted protein exhibits even higher similarity to F-spondin, we named the CG11739-gene product, Drospondin. Our data show that Drospondin is an extracellular matrix (ECM) protein secreted by glial cells. When added to cell cultures, Drospondin stimulated the arborization of MB neurons in a LpRs-dependent way. *In vivo*, the downregulation of Drospondin caused MB developmental defects through a genetic interaction with LpRs. Moreover, flies deficient in Drospondin exhibited alterations in sleep homeostasis, locomotion, and social behavior. These flies also showed a significant decrease in brain size. The structural defects in the brain and cultured MBs neurons were rescued when human Reelin was expressed under the control of the Drospondin promoter in Drospondin-deficient flies. These findings are consistent with Drospondin being a functional ortholog of vertebrate Reelin that controls the development and function of the *Drosophila* nervous system.

## RESULTS

### Drospondin is an extracellular glycoprotein with sequence similarity to F-spondin and Reelin

To identify the most similar gene to human *RELN* in *Drosophila* genome, we used the DIOPT-Ortholog prediction tool [26]. We also searched for *Drosophila* proteins containing the highest sequence homology to each of the Reelin domains, including the Reeler domain and the so-called Reelin repeats (Figure 1). An uncharacterized protein encoded by the *CG17739* gene emerged as the main candidate. This protein exhibits 29% amino acid sequence similarity with human Reelin (Supplementary Table S1). Our studies showed that CG17739-encoded protein was also similar to human F-spondin, with a 47% amino acid sequence similarity (Supplementary Table S1). Like Reelin, F-Spondin (also known as Spon1), is an ECM protein that binds to ApoER2 and VLDLR, regulating neuronal migration across developing cortical layers such as the floor plate of the neural tube, the hippocampus, cerebral cortex and the olfactory bulb [27–29].

**Figure 1.**
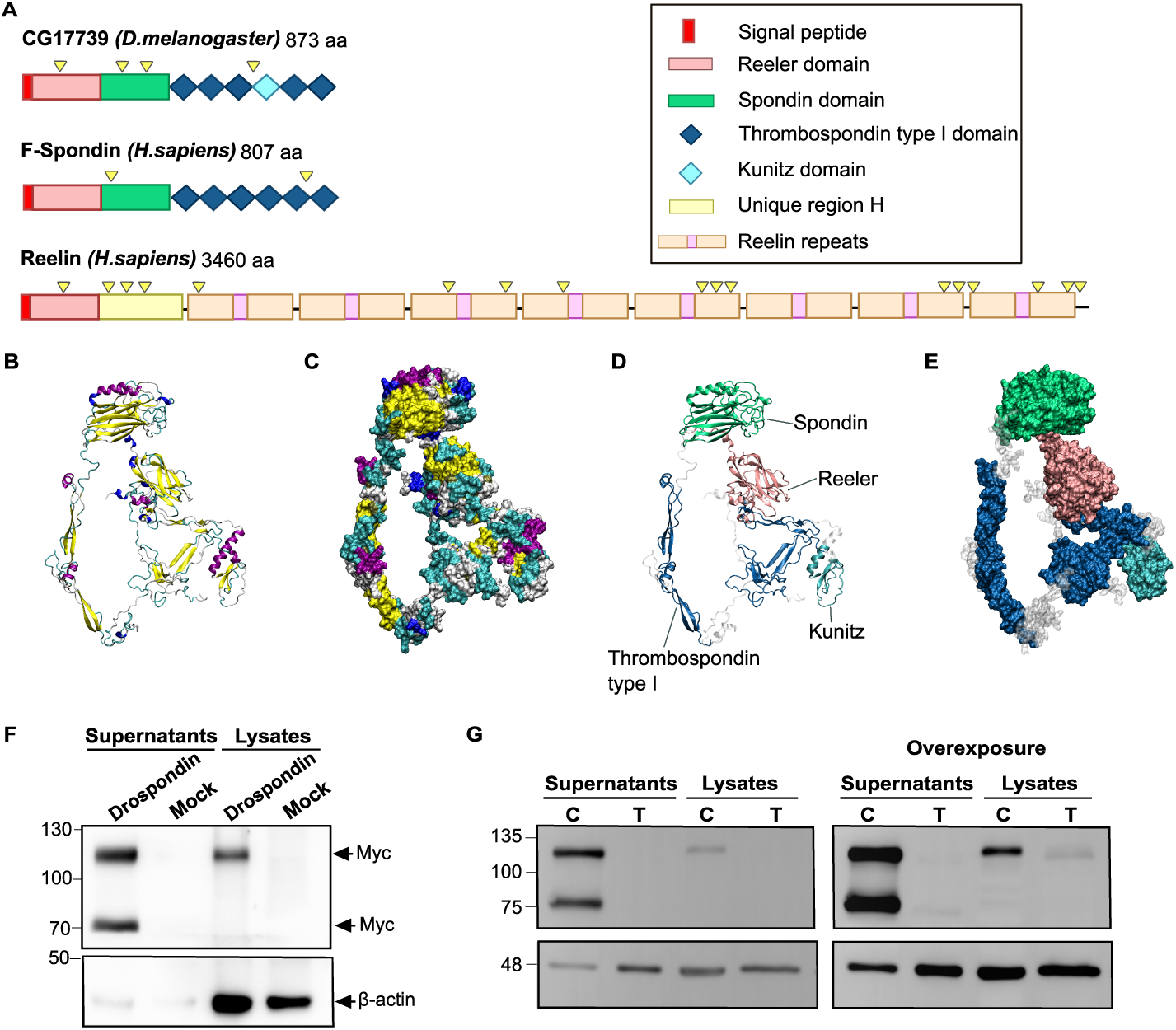
CG17739 protein is a glycoprotein with homology to mammalian F-Spondin and Reelin. (A) Schematic representation of CG17739-encoded protein, F-Spondin and Reelin with their different functional domains. N-glycosylation sites are indicated by yellow arrowhead. (B-E) Drosophila CG17739-encoded protein structure predicted via the RoseTTAfold method (Robetta server). Representations in cartoon (B and D), and in molecular surface (C and E) formats, were created with the VMD (Visual Molecular Dynamics) software. In (B) and (C) colors show the secondary structures (yellow: β-sheets, purple: α-helix, blue: coils, white: no defined structure), while in (D) and (E) different colors show CG17739 representative domains (red: reeler domain; green: spondin domain; blue: thrombondospin type I domain; light blue: Kunitz domain). (F) Western blots of Drospondin (CG17739-encoded protein) expression in S2 cells lysates and their supernatants. Two bands are detected in the supernatants; n=2 independent experiments. (G) Western Blot of S2 cells expressing CG17739 treated with Tunicamycin and their supernatants. Only after overexposure of the membrane it is possible to detect the band at a lower molecular weight. “C” and “T” indicate control and tunicamycin treatment, respectively; n=2 independent experiments.

The *CG17739* gene is predicted to produce only one transcript of 2,830 nucleotides (flybase.org, FBgn0033710) and an ECM protein of 873 aminoacids and 98,282 kDa (uniprot.org, Q7K3Y9_DROME). We generated a model for the organization of domains in the CG17739-encoded protein, based on data available in the UniProt server. This model was compared to the domain organization of human Reelin and F-spondin (Figure 1A). As expected for secreted proteins, all contain an N-terminal signal peptide that allows their translation and maturation at the rough endoplasmic reticulum. Additionally, the CG17739-encoded protein contains one Reeler domain, followed by a Spondin domain and Thrombospondin type I domain repeats (TSRs). In the case of Reelin, after the Reeler domain, there is a unique region followed by Reelin repeats. Drospondin also comprises a Kunitz domain, absent in F-spondin and Reelin. We generated a three-dimensional model of Drospondin using the RoseTTAFold method (Robetta server) [30] (Figures 1B-1E). The Reeler and Spondin domains exhibit β-sandwich conformations and are physically close to each other (Figures 1B and 1D). The TSRs exhibit antiparallel β-strand conformations, while the Kunitz domain has the characteristic combination of α helix and β strand (alpha-beta fold) (Figures 1B and 1D). Given all the similarities between the vertebrate and the CG17739-encoded protein, we named this protein Drospondin.

To corroborate that the identified *Drospondin* open reading frame (ORF) encodes a secreted protein, we cloned and expressed it in *Drosophila* S2 cells (Figure S1A). Drospondin was tagged with a N-terminal Myc epitope (Figure 1F, Figures S1A-S1C). Immunoblot analysis for the Myc epitope detected two bands of ∼85 and ∼120 kDa in the culture medium. Interestingly, only the ∼116.7-kDa band is detected in the cell lysates (Figure 1F; Figures S1B-S1C). This supports that the higher band is the full-length protein while the lower band may arise from cleavage of the larger species right before or after secretion into the extracellular medium. The 120 kDa band corresponds to a protein with a molecular mass higher than that predicted from the amino acid sequence (Figure 1F). A search for glycosylation sites revealed the presence of both N-glycosylation and O-glycosylation motifs in the Drospondin sequence [31]. We treated S2 cells stably expressing Myc-Drospondin with 5 μg/mL tunicamycin to inhibit N-glycosylation. This treatment reduced the amount of Drospondin in both the medium and cell lysates (Figure 1G, Figure S1D). The tunicamycin treatment also resulted in a lower molecular mass band, which is only evident after membrane overexposure, consistent with the proposition that Drospondin is N-glycosylated (Figure 1G). These findings suggest that Drospondin N-glycosylation is needed for its correct folding, maturation, stability and secretion, as is the case for other glycoproteins [32, 33].

### Drospondin is expressed in glial cells

Analysis of published single-cell RNA-seq data of the adult *Drosophila* brain [34] show that Drospondin is mainly expressed in glial cells. To verify this, we used a fly strain in which the last two exons of Drospondin are replaced with a CRIMIC cassette [35, 36] (hereafter called *Drosp^CR70269^*). The CRIMIC cassette includes a *T2A-GAL4* element followed by a stop codon. This produces a severe loss of function allele while expressing the GAL4 transcription factor in the expression pattern of endogenous Drospondin. Thus, this genetic tool can be used for expression of any UAS-transgene. Drospondin-expressing cells were identified by nuclear expression of mCherry fluorescent protein using *UAS-mCherry-NLS* (Figure 2A). Drospondin was found in some Repo-positive glial cells at the larval 3 stage (L3), and in almost all Repo-positive glial cells in pupae and adult flies (Figure 2A). The expression pattern of Drospondin suggests that it is mainly found in the ensheathing and cortex glia [37, 38]. Further studies revealed no colocalization of Drospondin and the neuronal marker Elav at any of the analyzed developmental stages, indicating that the protein is not expressed by neuronal cells (Figure 2B).

**Figure 2.**
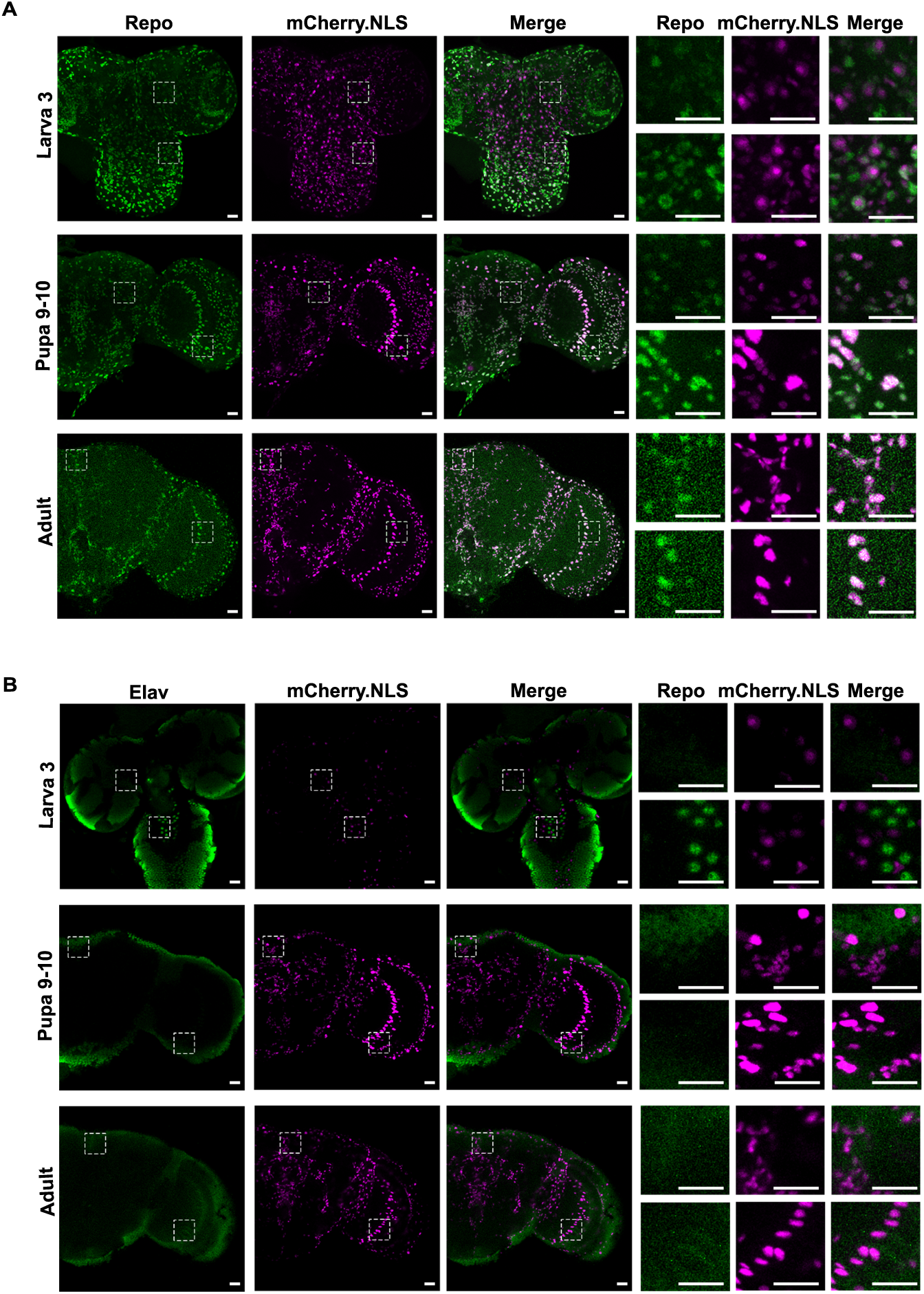
Drospondin is expressed by glial cells. (A) Representative microphotograph of immunostaining against Repo, a glial marker, in brains of flies expressing the fluorescent mCherry protein in the nucleus, directed by the Drospondin promoter (*Drosp^CR70269^*;+; *UAS*-*mCherry.NLS*). (B) Representative results of immunostaining against Elav in brains of flies expressing the fluorescent protein mCherry in the nucleus directed by the Drospondin promoter (*Drosp^CR70269^*;+; *UAS*-*mCherry.NLS*). Representative images of experiments carried out in L3 larvae, pupae (stage 9-10) and adult flies (0-3 days old), showing results for Repo or Elav staining, mCherry fluorescence and merged images. Three panels at right, show amplifications of areas indicated in panels at left. The white bars in the lower right corner of the images represents 20 µm.

### Drospondin increases the complexity of the MB neuronal arbor

Our previous work showed that treatment with mammalian Reelin increases the complexity of the neuritic arbor in cultured *Drosophila* MB neurons [17], similarly to the effects induced by Reelin in mammalian cortical and hippocampal neurons [18, 39, 40] or by F-spondin in mammalian neuronal cell lines [41]. We examined the role of Drospondin on MB neurons in culture, identified by expression of the fluorescent plasma membrane protein CD8-ChRFP under the control of the *c309-GAL4* driver (Figures 3A-3C). MB neurons were treated with recombinant Drospondin in the range of 0.5-50 nM, resulting in a higher complexity in the neuritic tree than controls (Figures 3A-3C). The maximum effect in the most extended neurite was found at a Drospondin concentration of 15 and 50 nM (Figure 3C). Thus, like Reelin, Drospondin increases the complexity of the neuritic tree of MB neurons in culture.

**Figure 3.**
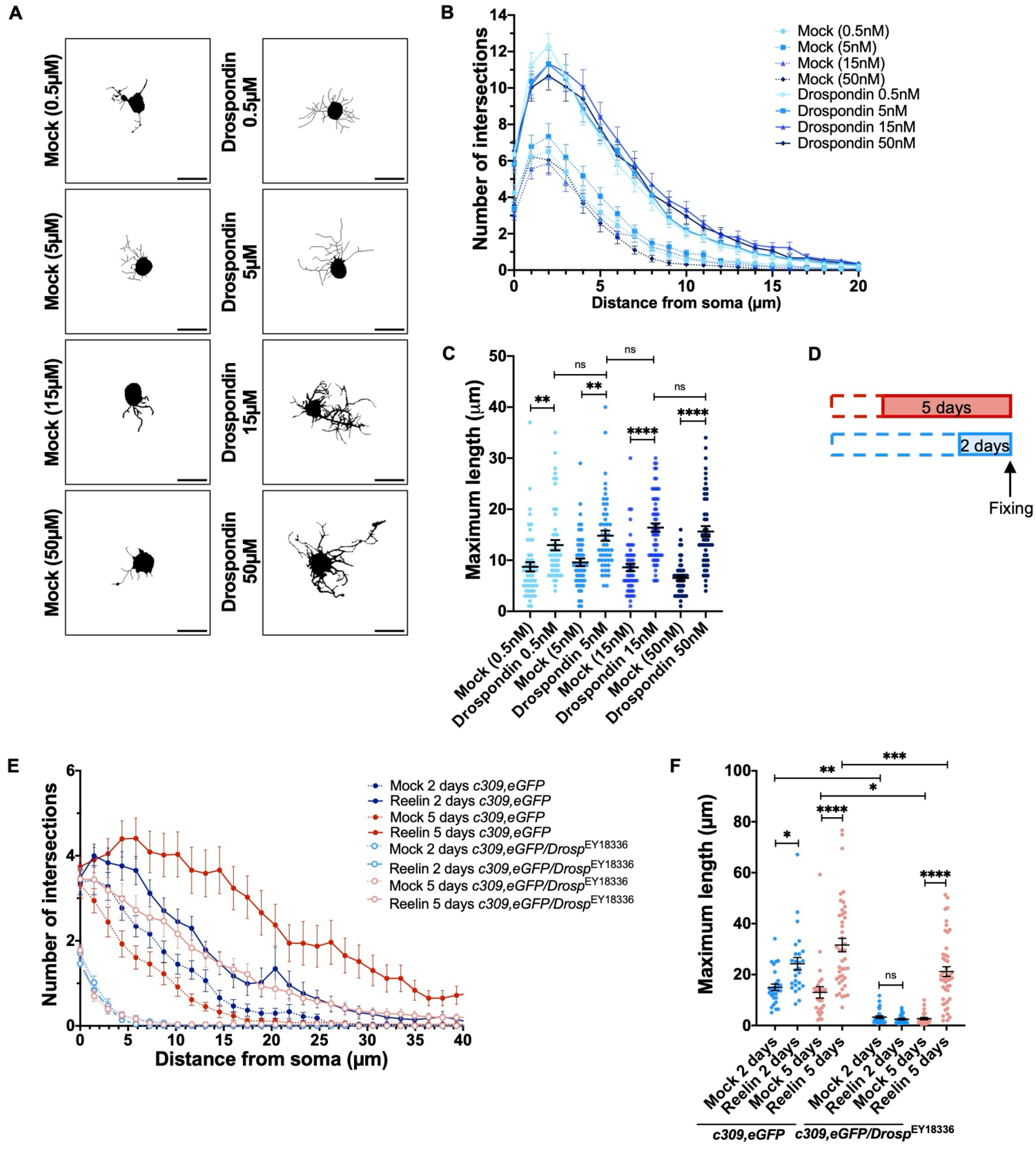
**Drospondin promotes neuritic growth in cultured neurons**. (A) Representative black and white images of primary neuronal cultures prepared from c309,CD8.ChRFP pupal brains, after treatment with Drospondin (at 0.5 nM, 5 nM, 15 nM and 50 nM concentrations) and their corresponding mock treatments. MB neurons were identified by RFP expression. The black bars in the lower right corner of each of the images represent 10 µm. (B) Sholl analysis of MB neurons under different experimental conditions. (C) Maximum length, a parameter defined as the larger distance reached by a neurite of a MB neuron with respect to its soma, under the different conditions assessed. Kruskal-Wallis test indicates overall p<0.0001 among all conditions. **, ****, indicate p<0.01, p<0.0001. “ns” not significant. Data from three independent experiments, each one consisting of 5 coverslips, 13-20 cells in each coverslip. Data are expressed as mean ± SEM. (D) Description of Reelin treatment. Primary brain cultures from Drospondin mutants were treated with Reelin for two or five days. Cells in culture were fixed at the seventh day *in vitro*, regardless the treatment. (E) Sholl profile of Drospondin mutants that express GFP only in MB neurons after Reelin treatment. Cells from Drospondin mutants are unable to respond to a 2-day Reelin treatment. However, when Reelin treatment is carried out for 5 days in the mutant neurons, their Sholl profile is similar to control cultures. (F) Neurite maximum length measured in neurons from Drospondin mutants treated with Reelin. Neurons from Drospondin mutants exhibit a decreased neurite length. A two-day Reelin treatment of Drospondin mutant cells does not affect their maximum length. However, when the Drospondin mutant cells are treated with Reelin earlier in their *in vitro* development (beginning by day 2 *in vitro*), the mutant cells recover their neurite maximum length to levels comparable to those observed in control cultures. In D-F, all cultures were prepared from haploinsufficient mutants for Drospondin. Data expressed as mean ± SEM, Two-way ANOVA, interaction p<0.05, Reelin treatment and genotype factors p<0.0001. Tukey post-test indicate *, **, *** and ****, which means p<0.05, p<0.01, p<0.001 and p<0.0001, respectively. n=3 coverslips from three independent experiments, 15-20 cells for each experimental serie.

We hypothesized that neurons of animals deficient in Drospondin expression might show a basal impairment in the development of their neuritic arbor and that Reelin might be able to rescue this deficit. To address these ideas, we studied the effect of Reelin on cultured MB neurons generated from Drospondin-deficient flies and that express cytoplasmic GFP (Figures 3D-3F). These experiments were performed using Drospondin heterozygous deficient animals containing an insertion in the 5’UTR of the gene (identified from here onwards as *Drosp^EY18336^*; Figures S2A and S2B). Primary cultured neurons were maintained for a total of seven days, and they were treated with recombinant Reelin for the last two or five days (Figure 3D). MB neurons from *Drosp^EY18336^* heterozygous mutants exhibited a basal deficit in neurite length and complexity compared to control neurons (Figures 3E and 3F). Importantly, these defects were not detected when the mutant neurons were treated with Reelin for 5 days: under these conditions, neurite arborization levels were comparable to those observed in the mock-treated control group. Reducing the Reelin treatment to the last two days of culture, however, showed no restoration of neurite growth (Figures 3E and 3F). Therefore, an early Reelin treatment rescued the deficit in neurite growth observed in MB neurons from Drospondin-haploinsufficient flies. These results suggest a critical time window for the role of Drospondin in neuronal differentiation, which can be substituted by Reelin.

Altogether, these findings indicate that Drospondin shares conserved functions with Reelin and F-spondin in facilitating neuritic outgrowth.

### Drospondin is required for MB axonal development

The MB is a structure in the *Drosophila* brain linked to several behaviors [21–23] exhibiting a particular anatomical organization where soma, dendritic and axonal segments are separated from each other; the axons form the so-called MB lobes [21, 42, 43]. Our previous work showed that flies with diminished expression of LpRs exhibit defects in MB development [17]. Since Drospondin could be a putative ligand for LpRs, we evaluated whether it localizes near the MB region. This was assessed using the *Drosp^CR70269^* allele to direct the expression of HA-tagged Drospondin under the control of the endogenous promoter (Figure 4A, Figure S3). FasII immunostaining was used to reveal the MB peduncles and lobes [44, 45]. Drospondin was mainly present on the surface of the larval brain without any evident localization in the MB. In pupae and adult animals, Drospondin was also present on the brain surface, as well as around the MB calix, peduncles and lobes. This staining pattern is consistent with a glial expression of Drospondin (Figure 4A) [37, 38, 46].

**Figure 4.**
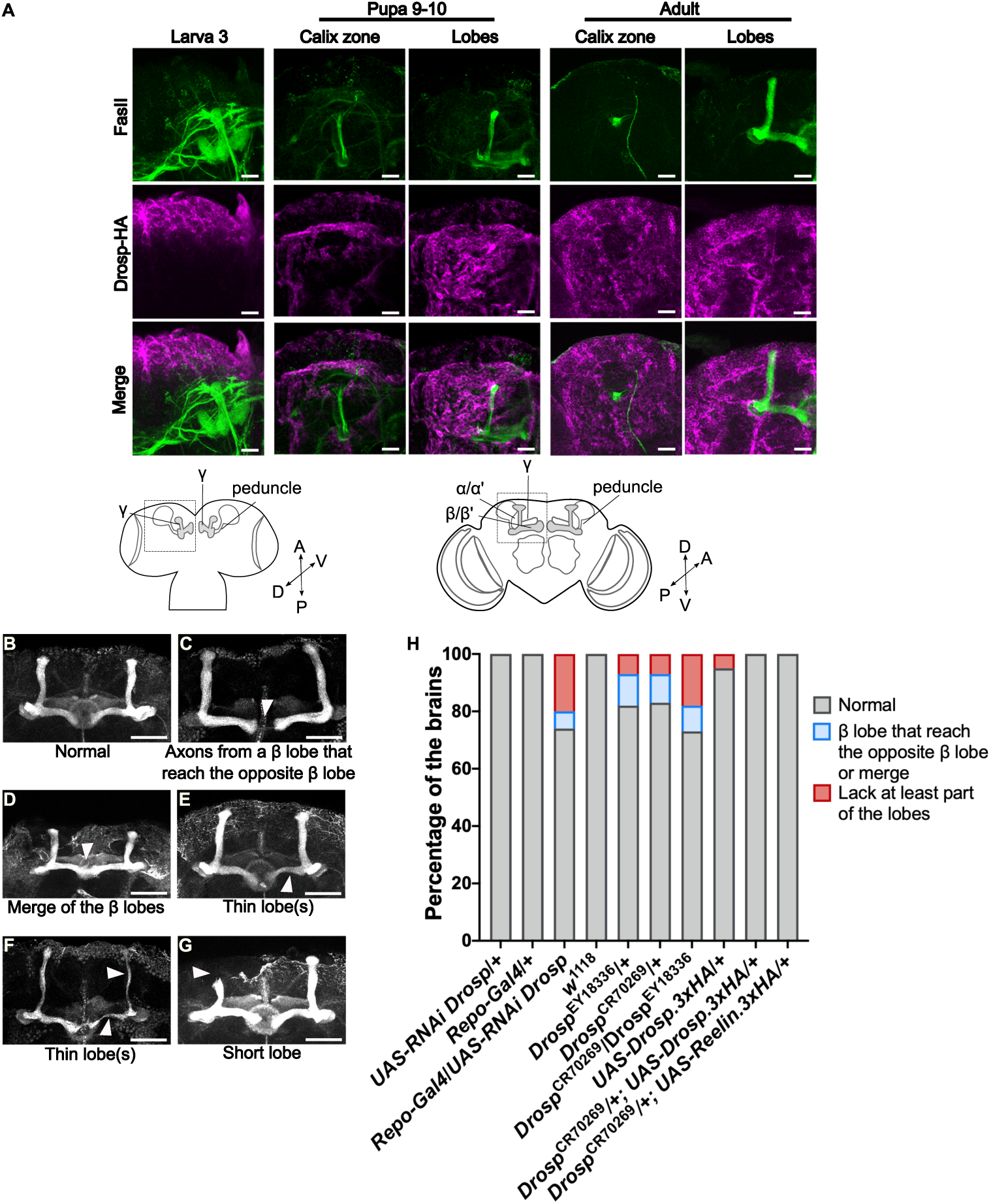
Drospondin contributes to MB adult structure. (A) Microphotographs of representative experiments showing the MB in larval, pupal and adult brains. It is shown the lobes and calyx regions in pupa and adult animals; peduncles, α and β lobes are identified by FasII immunostaining (green hue). Drospondin expression was evidenced by anti-HA staining (in magenta) in flies where *3HA-*tagged Drospondin was expressed under the control of endogenous promoter, by using the GAL4-UAS system (*Drosp^CR70269^*/+;*UAS-Drospondin.3xHA*). Data is representative of n=5, 7 and 17 L3 larvae, pupae and adult brains, respectively. The white bars at the lower right corner of each of the images represent 20µm. Bottom, scheme of larval and pupal/adult brains showing the regions studied. (B-G) Representative images of the phenotypes found with the different tools used to reduce Drospondin expression. (B) Typical MB structure in control conditions. (C) Axons from one β lobe that reach the opposite β lobe (*Repo-GAL4*/*UAS-RNAi Drosp*). (D) Merge of β lobes (*Drosp^CR70269^*/*Drosp^EY18336^*). (E) Thin lobe(s), in the microphotograph it is shown a thin β lobe (*Drosp^CR70269^*/*Drosp^EY18336^*). (F) Thin lobe(s), in the image α and β are thinner than normal (*Drosp^CR70269^*/+). (G) Short lobe, in the image a short α lobe is shown (*Drosp^CR70269^*/*Drosp^EY18336^*). The arrowhead indicates the abnormality. The white bars at the lower right corner of the images represent 50µm. (H) Percentage of fly brains from each genotype exhibiting identified phenotypes; data is presented per strain (in total number of brains assessed, number of independent experiments): *Repo-Gal4*/+ (25,2); *UAS-RNAi Drosp*/+ (23,2); *Repo-Gal4*/*UAS-RNAi Drosp* (35, 2); *w1118* control animals (36,3); *Drosp^EY18336^*/+ (24,2) *Drosp*^CR70269^/+ (30, 3); *Drosp^CR70269^*/*Drosp^EY18336^* (34,3); *UAS-Drosp.3xHA*/+ (19,2); *Drosp^CR70269^*/+; *UAS-Drosp.3xHA*/+ (23,2); *Drosp^CR70269^*/+;UAS-Reelin.3xHA/+ (17,2).

The α’ and β’ lobes of the MB develop towards the end of the L3 stage, while α and β lobes finish their maturation at the pupal stage [43]. Thus, by the beginning of adulthood the mature MB structure has been established [43]. Since our studies suggest that Drospondin is expressed in glial cells surrounding the MB, we asked whether a reduction in the glial expression of Drospondin has any consequences for MB development (Figures 4B-4H, Table S2). We decreased Drospondin expression in glial cells by using an *UAS-RNAi* transgene designed to target Drospondin transcripts under the control of the pan-glial driver *Repo-Gal4* (Figure S2C). FasII immunostaining was used to reveal the MB architecture in adult flies. In contrast to the typical MB structure in control flies (Figure 4B), we observed some degree of β lobes fusion in flies expressing RNAi targeting Drospondin in glial cells. For instance, in some brains, it was possible to detect axons from one β lobe in one side of the brain crossing the midline towards the opposite side (Figure 4C), while in other brains, there was a complete fusion of β lobes (Figure 4D). Other phenotypes include gross alterations of the MB structure, like the presence of one (Figure 4E) or two (Figure 4F) thinner than normal lobes, missing or shorter lobes (Figure 4G). Thus, about 6% of flies with glial Drospondin knockdown (KD) exhibit some degree of β lobes fusion and 20% of these brains lack parts of the MB lobes (Figure 4H).

To corroborate the contribution of Drospondin to the MB architecture with an additional tool, we used the CRIMIC strain. Finding homozygous *Drosp^CR70269^* mutants was challenging (only 2.5% and 1.6% of male and female animals, respectively), and they died prematurely with a median survival of 3 days (Figure S4). On the other hand, the transheterozygous mutants *Drosp^CR70269^*/*Drosp^EY18336^* were more frequently found, although their median survival was four days (Figure S4). As expected, the heterozygous and transheterozygous mutants expressed Drospondin at a significantly reduced level (Figures S2D and S2E). Thus, the MB architecture was evaluated in the two mutants in haploinsufficiency (*Drosp^CR70269^*/+ and *Drosp^EY18336^*/+) and transheterozygosity (*Drosp^CR70269^*/*Drosp^EY18336^*). Flies mutant for Drospondin in one allele displayed phenotypes related to the fusion of β lobes in 11% (*Droso^EY18336^/+*) and 10% (*Drosp^CR70269^*/+) of the analyzed brains (Figure 4). This is not different from what we observed in the transheterozygous mutants (*Drosp^CR70269^*/*Drosp^EY18336^*), with 8.8% of the brains showing fusion of β lobes. Interestingly, when assessing gross anatomy of the MB region including missing lobes, about 7% of *Drosp^EY18336^*/+ and *Drosp^CR70269^*/+ brains exhibited this phenotype. Of note, this phenotype increased in prevalence to 17.6% of brains in transheterozygous mutants.

We also tested whether rescuing the expression of Drospondin in the haploinsufficient mutant genetic background, can correct the MB defective phenotypes. Our results show that restoring Drospondin expression in the *Drosp^CR70269^* mutant background by using a HA-tagged protein (*Drosp^CR70269^*/+;*UAS-Drosp.3xHA*/+) prevented the MB phenotypes (Figure 4H, Table S2).

Considering the similarities in Reelin and Drospondin effects on MB neurons in primary culture, we also evaluated whether the *in vivo* replacement of Drospondin by human Reelin rescued the phenotype in the *Drosp^CR70269^* mutant genetic background (Figure 4H). Remarkably, the MB architecture of flies expressing human Reelin in Drospondin expression domain allele (*Drosp^CR70269^*/+;*UAS-Reelin.3xHA*/+) displayed a normal MB architecture (Figure 4H, Table S2), demonstrating that human Reelin can functionally replace fly Drospondin in this context.

Overall, these results demonstrated that Drospondin plays a role in MB development, and it is functionally orthologous to mammalian Reelin.

### Drospondin interacts genetically with LpRs

Flies haploinsufficient for LpRs exhibit impairment in MB organization [17], which is a similar situation to what is observed in Drospondin-deficient flies. We decided to assess whether there is a genetic interaction between Drospondin and LpR1 or LpR2. For this, flies heterozygous for a CRIMIC element insertion in LpRs, *LpR1^CR70219^* and *LpR2^CR70220^* were used. These mutants showed 16.6% and 10.4% of brains with β lobes fusion, respectively, similar to the proportion of *Drospondin* heterozygous mutant flies exhibiting this phenotype (Figures 5A-5G, Figure S5, Table S3). Remarkably, in adult flies bearing one mutant allele for *Drospondin* and one for the LpR1 gene (*Drosp^CR70269^*/+; *LpR1^CR70219^*/+), the frequency of flies with phenotypes associated with fusion of β lobes increased dramatically to 40.7%. Heterozygous mutant flies for *Drospondin* and LpR2 genes (*Drosp^CR70269^*/+; *LpR2^CR70220^*/+) showed a modest increase in the incidence of this phenotype, to 16.6% of the brains studied compared to *LpR2^CR70220^*heterozygous flies (Figure 5G, Table S3). We next assessed axonal growth defects in MB. About 5% and 8% of brains showed this phenotype in flies lacking one copy of LpR1 or LpR2, respectively (Figure 5G). Interestingly, the brains of *Drosp^CR70269^*/+; *LpR1^CR70219^*/+ flies showed almost no change in the frequency of defects, 7.4% of tissues show phenotypes. On the other hand, in the double mutants *Drosp^CR70269^*/+; *LpR2^CR70220^*/+, this phenotype was increased to 25% of the brains evaluated (Figure 5G). These results indicate that LpR1 and LpR2 genetically interact with Drospondin to modulate different aspects of MB development, supporting their participation in a common signaling pathway.

**Figure 5.**
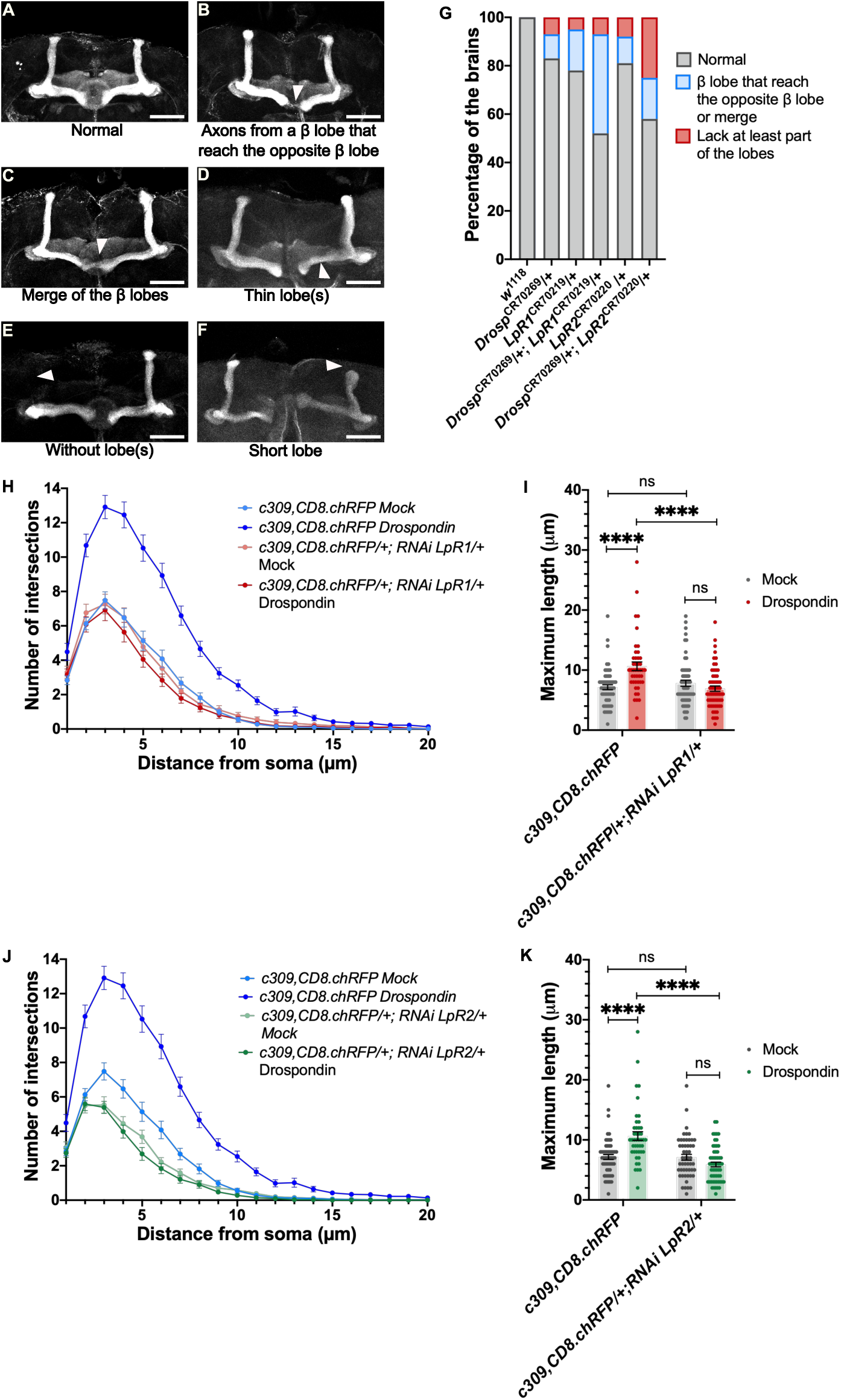
Drospondin and LpRs genetic interaction. Representative images for the immunostaining of MB using FasII in haploinsufficient flies for Drospondin and LpR1 or LpR2. (A) MB in control animals (*w^1118^);* (B) Axons from one β lobe that reach the opposite β lobe (*LpR2^CR70220^*/+). (C) merge of β lobes (*LpR1^CR70219^*/+). (D) Thin lobe(s), it is shown a thin β lobe (*Drosp^CR70269^/+; LpR1^CR70219^*/+). (E) Short lobe, in the image a short α lobe (*Drosp^CR70269^/+; LpR2^CR70220^*/+). (F) Without lobe(s), lack one α lobe (*LpR2^CR70220^*/+). The arrowhead indicates the abnormality. The white bars in the lower right corner of the images represent 50µm. (G) Distribution (in percentage) of each phenotype in animals of genotypes indicated. Data is presented per strain (in total number of brains studied, number of independent experiments): *w^1118^*control animals (14,2); *Drosp^CR70269^*/+ (30, 3) (same data used in Figure 4 panel H, experiments carried out in parallel)*; LpR1^CR70219^*/+ (60,4); *Drosp^CR70269^/+; LpR1^CR70219^*/+(27,2) *LpR2^CR70220^*/+ (48,3); *Drosp^CR70269^/+; LpR2^CR70220^*/+ (24,2). (H) Sholl analysis of primary neuronal cultures from control animals or flies with a downregulation for LpR1 expression in MB, exposed to Drospondin. (I) Maximum length recorded in MB neurons in each condition. Two-way ANOVA, interaction p<0.0001, genotype factor p<0.005 and treatment factor p<0.05. Tukey post-test, **** means p<0.0001, “ns” means not significant. (J) Sholl analysis of control MB neurons or neurons with a downregulation for LpR2 expression, after exposure to Drospondin. (K) Maximum length in MB neurons in each condition. Two-way ANOVA, interaction p<0.0001, genotype factor p<0.0001 and treatment factor p<0.05. Tukey post-test, **** means p<0.0001, “ns” means not significant. Data (in H-K) from 4 different coverslips in three independent experiments, 13-20 cells in each experiment. Data are expressed as mean ± SEM.

As the Reelin-induced response in cultured MB neurons depends on LpRs [17], and Drospondin promotes neuritic tree outgrowth (Figures 3A-C), we then asked whether the Drospondin-induced response in cultured MB neurons depends on LpRs. The MB neurons KD for LpR1 (*c309,CD8.chRFP*/+; *RNAi LpR1*/+) were unable to respond to 15 nM of Drospondin by incrementing neither the complexity of their neuritic arborization (Figure 5H), nor the maximum length that their neurites can reach (Figure 5I). MB LpR2 KD neurons (*c309,CD8.chRFP*/+; *RNAi LpR2*/+) also failed to respond to the Drospondin treatment (Figures 5J-5K).

Altogether, these results show that LpRs genetically interact with Drospondin to regulate MB development and neuritic tree complexity.

### *Drospondin*-downregulated flies exhibit smaller brain size

Mutations in *RELN* have been linked to disorders like lissencephaly with microcephaly and cerebellar hypoplasia [9, 47, 48]. These alterations are thought to arise from decreased Reelin signaling and abnormal whole-brain development [47, 48]. The expression pattern of Drospondin indicates that this protein is expressed in glial cells distributed throughout the brain, implying that several cell types and brain regions could receive and be influenced by this signaling molecule.

To evaluate whether flies with decreased Drospondin expression exhibit any significant whole-brain alteration (Figure 6) we measured the middle brain area (Figure 6A) of animals with Drospondin downregulation only in glial cells (*Repo-GAL4*/*UAS-RNAi Drosp*). Remarkably, these flies exhibited a smaller brain area (Figure 6D) as compared to genetic controls (Figures 6B-6C) (*Repo-GAL4*/+ and *UAS-RNAi Drosp*/+). It is noticeable that 19% of the brains in the KD experimental group were smaller than the smallest brains recorded in any of the two genetic controls (red bar in Figure 6E). Since severe microcephaly is defined in humans by a birth head circumference that is three or more standard deviations (SD) below the mean average size in the entire population [49–51], we quantified the number of brains that exhibit a central brain area smaller than three SD the mean average recorded in control genotypes. Our data show that 33% of the brains of flies with Drospondin KD in glial cells fulfil the criteria for microcephaly.

**Figure 6.**
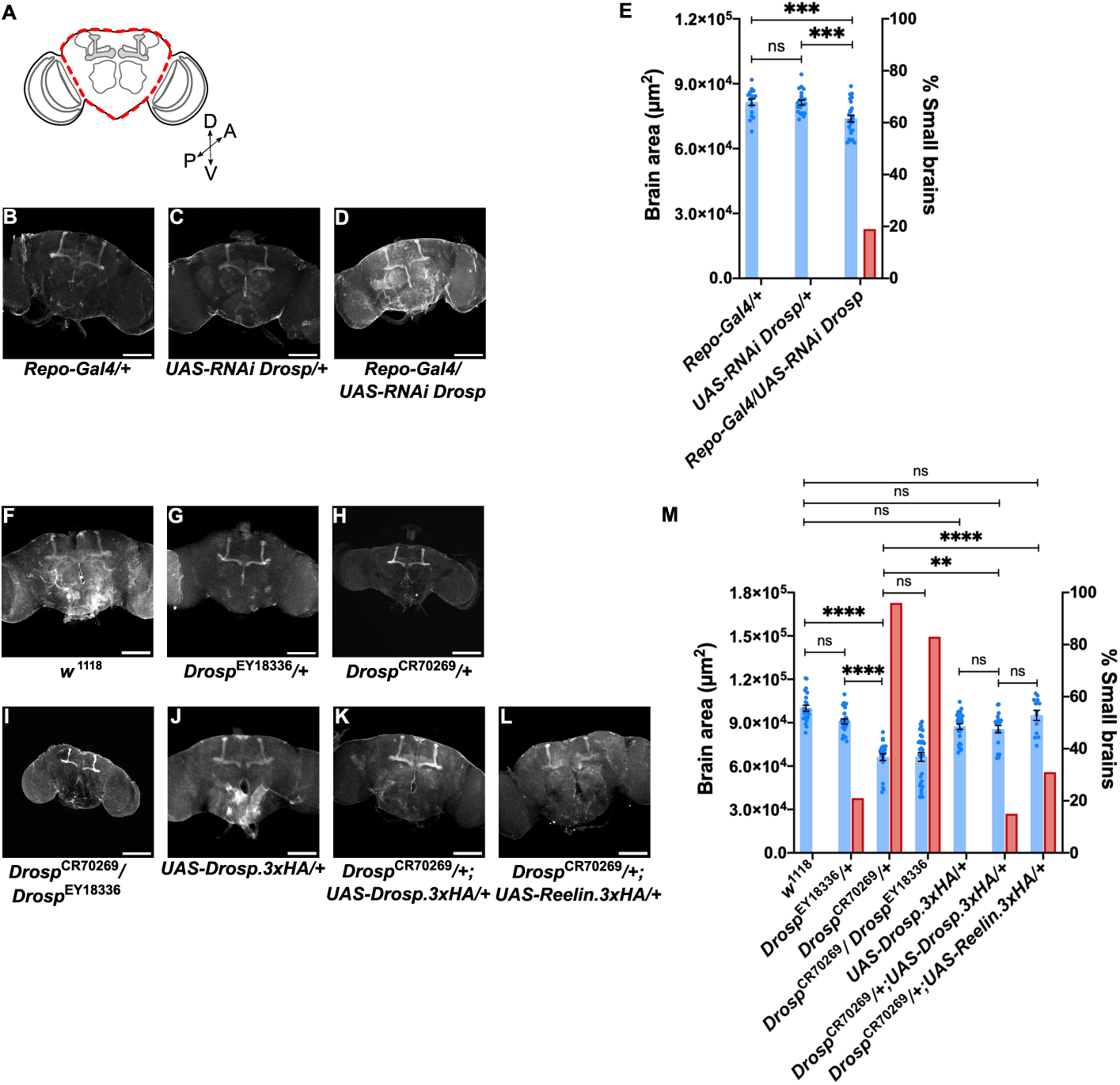
**Drospondin missexpression affects brain size. (**A) Scheme of the brain area measured after the Z-axis projection. The discontinuous red line represents the measured area (middle brain). (B-D) Representative brain images obtained in control strains (B) *Repo-GAL4*/+ and (C) *UAS-RNAi Drosp*/+, and (D) from flies with knockdown for Drospondin (*Repo-GAL4*/*UAS-RNAi Drosp*). (E) Quantification of middle brain area in flies from each genotype. One way-ANOVA indicated p<0.0001 over all conditions; Tukey posteriori test show *** which indicates p<0.001 in experimental group as compared to control strains. “ns” means not significant. Right y-axis shows percentage of brains with a smaller area than the smallest brain area recorded in the two controls (in red). Data is presented per strain (in total number of brains, number of independent experiments): *Repo-GAL4*/+ (17,2); *UAS-RNAi Drosp*/+ (21,2); *Repo-GAL4*/*UAS-RNAi Drosp (26,2).* (F-L) Representative images of the middle brain area from (F) control animals (*w^1118^*); (G-H) heterozygous mutant flies for Drospondin (*Drosp^EY18336^*/+ and *Drosp*^CR70269^/+); (I) transheterozygous mutant flies for Drospondin (*Drosp^CR70269^*/*Drosp^EY18336^*); (J) control flies bearing the undriven genetic element to express Drospondin under control of UAS (*UAS-Drosp.3xHA*/+), (K) animals expressing Drospondin under the control of the endogenous promoter, in the Drospondin CRIMIC genetic background (*UAS-Drosp.3xHA/+; Drosp^CR70269^/+*), (L) flies expressing Reelin directed by Drospondin promoter in the Drospondin CRIMIC background (*UAS-Reelin.3xHA/+; Drosp^CR70269^/+*). (M) Middle brain area recorded in each genotype. Kruskal-Wallis test p<0.0001 followed by Dunn post-test. ** and **** indicate p<0.01 and p<0.0001, respectively. “ns” means not significant. Right y-axis show percentage of brains with a smaller area than the smallest brain area recorded in either of the controls, *w^1118^* or *UAS-Drosp.3xHA*/+ (in red). Data is presented per strain (in total number of brains, number of independent experiments): *w^1118^* (23,3); *Drosp^CR70269^*/+ (23, 3); *Drosp^EY18336^*/+ (24,2); *Drosp*^CR70269^/*Drosp^EY18336^* (29,3); UAS-Drosp.3xHA/+ (27, 2); *Drosp^CR70269^/+;UAS-Drosp.3xHA/+* (20, 2); *Drosp^CR70269^/+; UAS-Reelin.3xHA/+* (13, 2).

We also studied this phenotype in Drospondin mutant animals (Figures 6F-6M). Although about 21% of the *Droso^EY18336^*/+ mutants exhibited a brain area smaller than the smallest area recorded in control animals (red bars in Figure 6M), the mean brain area measured in *Droso^EY18336^*/+ mutant flies was not significantly different than that recorded in the control strain (Figures 6F, 6G and 6M). In contrast, a significantly smaller mean brain area was measured in *Drosp^CR70269^*/+ as compared to controls (Figure 6F, 6H and 6M). In the heterozygous CRIMIC mutants, 83% of the brains had a brain area smaller than that recorded in control flies (red bar, Figure 6M), and 59% percent of them fulfilled the criterion for microcephaly. The differential effects between the two mutants could be explained by the level at which each genetic allele decreases Drospondin expression, with the CRIMIC mutant being more potent (Figure S2A, S2B and S2E). The transheterozygous mutant (*Drosp^CR70269^*/*Drosp^EY18336^*) also showed a significantly smaller brain area as compared to control flies (Figures 6F, 6I and 6M), which was not different from the effect detected in the heterozygous CRIMIC mutant (Figure 6M). Importantly, the expression of Drospondin driven by the CRIMIC allele (*Drosp^CR70269^*/+; *UAS-Drosp.3xHA*/+) resulted in a full rescue of the brain size, since the averaged brain area recorded is not different from its genetic control (*UAS-Drosp.3xHA*/+) or the *w^1118^* strain (Figures 6F, 6J, 6K and 6M). On the other hand, the expression of Reelin in the *Drosp^CR70269^* mutant background (*Drosp^CR70269^*/+;*UAS-Reelin.3xHA*/+) was also efficient at rescuing the brain size phenotype (Figures 6F, 6L and 6M), although it is still possible to detect about 35% of smaller brains.

Taken together, these results support the notion that Drospondin is a relevant glial signaling molecule in *Drosophila* whole-brain development.

### Drospondin deficiency throughout development impacts sleep, locomotion and social distance in adult animals

As mentioned, the MB region is associated with the regulation of sleep homeostasis and locomotion in flies. Given the structural phenotypes observed in the MB of Drospondin mutants (Figure 4), we asked whether *Drosp^CR70269^*/+ animals exhibited behavioral alterations. First, we assessed circadian motor activity when flies are under a 12:12 h light:dark cycle (Figure 7A-7D). The *Drosp^CR70269^*/+ mutant exhibited an overall longer sleeping time primarily due to a defect during the light phase, as there were no differences during the dark phase (Figure 7B). In contrast, during the light phase, there was a shortened latency to sleep in mutant animals as compared to controls (Figure 7C). Drospondin heterozygous mutants also presented an increased number of sleep events (Figure 7D). Thus, *Drosp^CR70269^*/+ mutants exhibited a dysregulation in the circadian sleep pattern.

**Figure 7.**
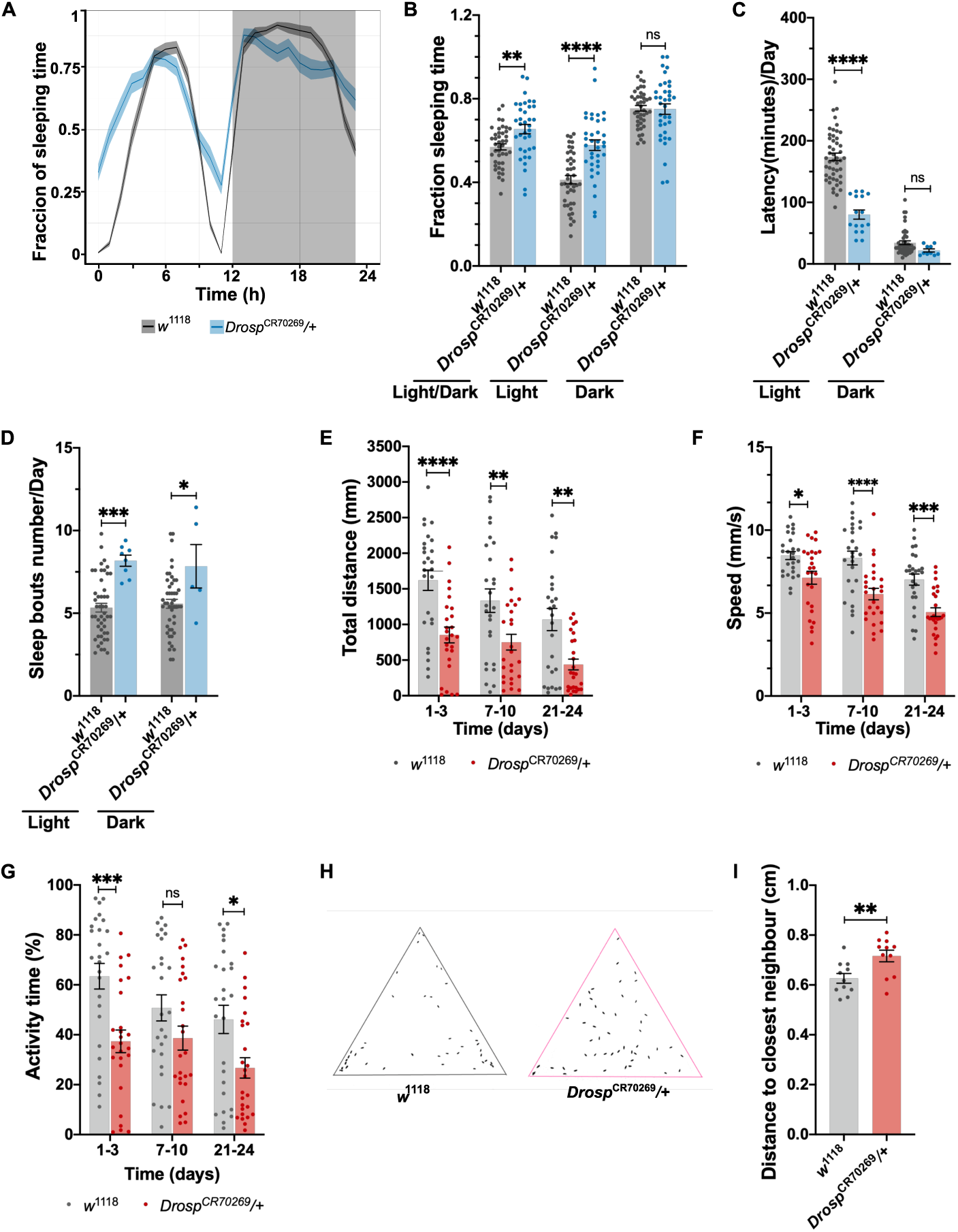
Impairment in sleep pattern, locomotor activity and social distance in mutants for Drospondin. (A) Sleep profile of haploinsufficient mutant flies for *Drospondin* (*Drosp^CR70269^*/+) as compared to control flies (*w^1118^*). The gray shadowed background represents the hours when flies were at dark. (B) Fraction of sleeping time recorded in the two fly strains, in light and dark conditions. Two-way ANOVA, interaction of factors p<0.0005, light-dark factor p<0.0001 and genotype factor p<0.0001. Bonferroni post-test indicates ** and ****, representing p<0.01 and p<0.0001 respectively. ns, not significant differences in dark conditions. (C) Average latency per day. Two-way ANOVA, p<0.0001 for interaction of factors, genotype and light-dark factor. Bonferroni post-test indicates ****, indicating p<0.0001. ns, not significant differences in dark conditions. (D) Average of number of sleep bouts per day. Two-way ANOVA; interaction of factors p=0.6114, light-dark factor p=0.9305 and genotype factor p<0.0001. Bonferroni post-test indicates * and ***, with p<0.05 and p<0.005, respectively. Data (in A-D) is presented per strain (number of animals, number of independent experiments): *w^1118^* control animals (42,3); *Drosp^CR70269^*/+ (36, 3). (E-G) Locomotor activity recorded in adult flies haploinsufficient for Drospondin (*Drosp^CR70269^*/+) as compared to control animals (*w^1118^*), at different time windows. (E) Total distance traveled by flies of the two strains at the different time windows. Two-way ANOVA; interaction of factors p=0.7646, time factor p<0.005 and genotype factor p<0.0001. Bonferroni’s multiple comparison test show **, **** which means p<0.01 and p<0.0001 respectively. (F) Average speed per fly in the times studied. Two-way ANOVA; interaction of factors p=0.4415, time factor p<0.0001 and genotype factor p<0.0001. Bonferroni’s multiple comparison test indicate *, ***, **** meaning p<0.05, p<0.005 and p<0.0001 respectively. (G) Activity time, represents the percentage of time when each fly was moving. Two-way ANOVA; interaction of factors p=0.3701, time factor p<0.05 and genotype factor p<0.0001. Bonferroni’s multiple comparison test, *, ***, means p<0.05 and p<0.005 respectively, “ns” means not significant. n=26 animals per genotype at each time window. (H-I) Social distance recorded in a triangular arena for heterozygous flies, mutants for Drospondin (*Drosp^CR70269^*/+) and control flies (*w^1118^*). (H) Representative images of experiments where flies of the two genotypes were allowed to position themselves in the triangular arena. (I) Averaged distance to the closest neighbor recorded in flies of the two genotypes in independent experiments. Unpaired t-test p<0.01. n=11 independent experiments, 30-45 flies per genotype in each experiment. Data is expressed as mean ± SEM in all the graphs.

Adult flies were also assessed in their basal locomotion at different ages (Figures 7E-7G). One-to three-day-old Drospondin mutants showed reduced walking distance (Figure 7E), speed (Figure 7F) and activity time (Figure 7G), as compared to controls, when placed in an open arena. In the 7-10-day-old window, the mutants showed reduced total distance traveled with no significant differences in the other motor parameters (Figures 7E-7G). In contrast, older flies showed a reduction in all motor parameters as compared to control flies. These results suggest that defects in brain development caused by Drospondin loss contribute to locomotor phenotypes in flies.

In humans, many neurodevelopmental disorders are associated with alterations in social behaviors [52–55]. As Reelin mutations are present in neurodevelopmental disorders, including some forms of ASD, characterized by a diminished ability to establish adequate social interactions [56], and the MB participates in social behaviors in flies [57], we evaluated social interaction in flies with reduced expression of Drospondin (Figures 7H and 7I). Drospondin-deficient animals maintained a higher distance to the closest neighbor (Figures 7H and 7I), a proxy for asocial behavior in flies [58, 59]. These data suggest that Drospondin plays a role in social behavior in *Drosophila*.

Overall, these results support that reduced expression of Drospondin throughout development has significant consequences for different behaviors associated with the MB.

## DISCUSSION

In the search for a *Drosophila* Reelin ortholog that explains the role of LpRs in MB formation and function, as well as the LpR-dependent Reelin-induced effect in cultured MB neurons [17], we identified the previously uncharacterized *CG17739* gene. We found that this gene encodes for an extracellular glycoprotein secreted by *Drosophila* glial cells in the brain, which we named Drospondin. Similar to Reelin effects on vertebrate [15, 18, 39] and invertebrate [17] neurons, cultured MB neurons respond to the addition of recombinant Drospondin by increasing the complexity of their neuritic tree in a LpRs-dependent manner. In addition, flies with reduced expression of Drospondin exhibit MB developmental defects, like those found in LpR deficient animals. Furthermore, Drospondin and LpRs genetically interact to influence MB development. Flies deficient in Drospondin throughout development also exhibit impaired sleep regulation, locomotion, and social distance. We also report that Drospondin expression is relevant for normal brain size in flies. Remarkably, vertebrate Reelin was able to rescue the neurite defects in MB neurons from Drospondin-deficient flies in primary cell culture, as well as the MB structure and brain size in adult flies. Therefore, Drospondin is a protein that was previously overlooked but has an essential role in brain development, the impairment of which has profound functional consequences for the animal. Overall, these findings are consistent with the notion that Drospondin might signal through a pathway analogous to that described for Reelin in vertebrates.

Our bioinformatics analysis indicates that Drospondin is more similar to F-spondin than to Reelin, consistent with previous amino acid sequence homology studies on members of the F-spondin family [60, 61]. It has been proposed that there is at least one other ortholog for vertebrate F-spondin in the *Drosophila* genome, encoded by *CG6953* and named fat-spondin [28]. Regarding their structures, vertebrate F-spondin and Reelin, as well as Drospondin and fat-spondin, contain a Reeler domain following the signal peptide. In addition to the Reeler domain, Drospondin, fat-spondin and F-spondin share the spondin (N-terminal) domain followed by TSR domains. All these domains are in the N-terminal region, which exhibit the highest sequence homology among them [41]. Fat-spondin and Drospondin are predicted to contain a Kunitz domain in the C-terminal half of the protein, which is not found in the vertebrate proteins [61]. The common structural features support the idea that these proteins could contribute to similar functions in vertebrates and *Drosophila*. Importantly, besides the description of the existence of fat-spondin, there are no reports on the contribution of this protein to any biological function in *Drosophila*.

### Drospondin contributes to brain formation

Literature indicates that in vertebrates, Reelin and F-spondin have similar roles but also some differences. For instance, F-spondin plays a role during floor plate development in vertebrates, allowing the pathfinding of commissural axons [27, 29, 61]. F-spondin mRNA is also present in embryonic Schwann cells when sensory and motor axons project to their peripheral targets [62]. To accomplish its actions, F-spondin binds to ApoER2 and VLDLR through their TSR domains [63–65], which Drospondin also contains. On the other hand, Reelin binds its receptors, ApoER2 and VLDLR, through their central tandem repeats to coordinate neuronal positioning and axon determination and growth [3, 66]. Regardless of the mechanism used, we proposed that Reelin, F-spondin and Drospondin contribute to neurite formation, axon pathfinding and brain development. Thus, we asked whether Drospondin contributes to fly brain development, focusing on a well-known structure, the *Drosophila* MB.

Drospondin-deficient animals exhibit a variety of anatomical defects. Some of the phenotypes were similar to those reported in flies deficient in LpRs including the fusion of β lobes [17]. This phenotype can be explained by axons crossing the midline of the brain. Since glial cells expressing Drospondin surround the MB, we propose that this protein may act as a stop signal for axon extension. The demonstration of a genetic interaction between Drospondin and LpRs in modifying β lobe fusion, further supports that Drospondin would act as the ligand and LpRs as the receptors. This is consistent with a study in *C. elegans* reporting that the central TSR1-4 of F-spondin binds a Lipoprotein Receptor Related Protein (LRP), resulting in axon repulsion [67]. Likewise, Reelin promotes detachment of neurons from the radial glia once they reach layer V of the cerebral cortex and stop their migration in rodents [3, 68]. In addition, high concentrations of F-spondin induce the inhibition of neurite outgrowth [41]. Thus, our results suggest that Drospondin shares conserved functions with mammalian Reelin and F-spondin, an idea that finds support in the fact that vertebrate Reelin reverts MB developmental defects in animals deficient in Drospondin.

Drospondin promotes the extension of *Drosophila* MB neurons processes since Drospondin deficiency results in thin or missing MB lobes, which could be attributed to either a reduction in the number of axons protruding from MB neurons or to axons failing to reach their targets. Primary neuronal cultures deficient in Drospondin exhibit poor neurite development, like the phenotypes of LpR-deficient animals [17]. Schubert et al., 2006 [41] previously reported that F-spondin promotes neurite outgrowth of nerve precursor cells in rats. Likewise, Reelin induces neurite development in hippocampal and cortical vertebrate neurons [15, 39, 40]. Importantly, our work shows that human Reelin can replace fly Drospondin during MB development, demonstrating a conserved function of the protein. Thus, our results support that fly Drospondin, mammalian Reelin and F-spondin, all similarly promote neuronal outgrowth and differentiation.

The Reelin-induced responses in *Drosophila* neurons depend not only on LpRs but also Dab, the fly homolog of mammalian Dab1 [17]. Dab1 is the first mediator of the canonical signaling cascade induced by Reelin in mammalian cells [3, 69, 70]. Similarly, the binding of F-spondin to ApoER2 triggers Dab1 phosphorylation [71, 72]. Therefore, a possible conservation in the signaling pathways between Reelin, F-spondin and Drospondin, is an attractive idea that needs to be assessed in the future.

Our study focused on the role of Drospondin in MB anatomy and functions. However, other brain areas could be affected by the lack of this protein since it is broadly expressed by glial cells throughout the entire brain. Indeed, we found a smaller brain size (microcephaly) of Drospondin-KD flies that could be explained by a smaller number of brain cells because of a delay in brain development, cell cycle defects, increased apoptosis in neuronal progenitors, defects in pattern formation, or lack of cellular differentiation [73, 74]. Further studies will be needed to explore these possibilities. Importantly, *RELN* and *VLDLR* mutations have been linked to lissencephaly with cerebellar hypoplasia in humans [9, 47, 48, 75]. Consistent with this evidence, changes in the size of different brain areas are found in the mouse mutant for the Reelin gene (known as the *Reeler* mouse), including a decreased size of the cerebellum and olfactory bulb [76–78]. Thus, the Drospondin mutant provides a new genetic tool to study cellular mechanisms of microcephaly.

### Drospondin contributes to *Drosophila* behavior

Reelin deficiency produces significant morphologic changes in the vertebrate brain. Further, Reelin loss-of-function mutations are associated with different neurodevelopmental and psychiatric disorders including ASD, schizophrenia, bipolar and mood disorders [10, 79–82]. Many of these conditions have been associated with alterations in sleep regulation, motor output and social interaction, among other features [83–86]. However, little is known on the direct contribution of Reelin deficiency to any of these behavioral features. One of the few reports available on this issue shows that patients with lissencephaly linked to *RELN* mutations exhibit abnormal neuromuscular connectivity [87] and hypotonia [47], which could underlie motor deficits. Less is known about a potential contribution of F-spondin genetic deficits to behavioral alterations in human disorders. In this regard, F-spondin is detected in the neocortex and hippocampus in adult animals, and it has been suggested that it may play a role in AD etiology by binding to ApoER2 and the amyloid precursor protein (APP) [61]. *Drosophila* constitutes a tool to gain a better understanding of the effects induced by a deficit in these signaling molecules, how these signaling molecules are linked to human disorders, and what are the molecular mechanisms involved in them.

The homozygous recessive *Reeler* mice exhibit dystonia, ataxia and tremor, while mice with one mutant copy of *Reln* are normal for these phenotypes [88]. On the other hand, it has been reported that mice lacking F-spondin lose their ability to maintain intrinsic circadian rhythmicity [89]. Accordingly, flies deficient in Drospondin showed alterations in the maintenance of homeostatic sleep, the execution of motor programs, and the regulation of social space, a proxy for social interactions. These are behavioral phenotypes previously reported in *Drosophila* models for neurodevelopmental and/or mental disorders and are proof that fly models can inform on pathological mechanisms underlying related human conditions, as has been reported elsewhere [58]. Future studies should evaluate whether and to what extent specific brain areas and neuronal circuits underlying circadian or social behaviors are affected in Drospondin mutant flies.

In conclusion, our study has identified and characterized a novel signaling molecule in *Drosophila*, which we have named Drospondin. This extracellular glycosylated protein plays a key role in the development of the MB, uncovering a novel signaling pathway relevant to brain formation and function. Furthermore, our data support the notion that roles of *Drosophila* Drospondin are similar to the functions of mammalian F-spondin and Reelin. This work opens new research avenues into how brains attain their mature form to operate adequately and to sustain normal behaviors. Given the similarities between Drospondin and vertebrate Reelin and F-Spondin, this work also supports the use of *Drosophila* as an animal model to gain a better understanding of how the brain is affected by deficiencies in these signaling molecules. Such insights could shed light on the involvement of these molecules in neurodevelopmental disorders like ASD and schizophrenia, and to neurodegenerative diseases like AD or Parkinson’s Disease.

## MATERIALS AND METHODS

All procedures were approved by Comité de Ética y Seguridad, Pontificia Universidad Católica de Chile (ID Protocols: 190806006 and 220620004)

### Drosophila stocks

Flies were raised on standard fly food at 25° and under a Dark:light cycle of 12:12 hrs. The flies employed were generated using the following strains from Bloomington *Drosophila* Stock Center (BDSC): *UAS*-*mCherry.NLS*: *w*;;P{w+mC=UAS-mCherry.NLS}3* (RRID:BDSC_38424); *Drosp*^EY18336^: *y^1^w^67c23^*;*P{y+mDint2w+mC=EPgy2}CG17739^EY18336^* (RRID:BDSC_17316); RNAi for Drospondin: *w*\*;*P{y+t7.7v+t1.8=TRiP.JF03198}attP2*/*TM3,Sb1* (isogenized to *w^1118^* genetic background from RRID:BDSC_53498); *Repo-GAL4: w^1118^*;;*P{w+m*=GAL4}repo*/*TM3,Sb1* (RRID:BDSC_7415); RNAi for LpR1: *w*;P{y+t7.7v+t1.8=TRiP.JF02551}attP2* (from RRID:BDSC_27249); *Drosp^CR70269^*:*y^1^w\**;*TI{GFP3xP3.cLa=CRIMIC.TG4.1}CG17739^CR70269-TG4.1^*/ *SM6a* (RRID:BDSC_93823); *LpR1^CR70219^*: *y1w*;;TI{GFP3xP3.cLa=CRIMIC.TG4.0}LpR1^CR70219-TG4.0^ (*RRID:BDSC_93804*); LpR2^CR70220^*: *y1w\**;;*TI{GFP3xP3.cLa=CRIMIC.TG4.0}LpR2^CR70220-TG4.0^*/*TM3, Sb1Ser1 (*RRID:BDSC_93805); RNAi for LpR2: *w*;P{y+t7.7 v+t1.8=TRiP.JF01627}attP2 LpR2* (from RRID:BDSC_31150). The *fly C309,CD8::ChRFP* was constructed from the flies *w\**;*P{w+mW.hs=GawB}c309* (RRID:BDSC_6906) and *w\**;*P{w+mC=UAS-mCD8.ChRFP}2* (RRID:BDSC_27391)

### Drosophila fly line development

pGW constructs were inserted into the VK00033 docking site using φC31-mediated transgenesis [90] to generate *UAS-Reelin-HA* (*y^1^w*;;PBac{UAS-Reelin-3xHA}VK00033/TM3,Sb1Ser1*) and *UAS-Drosp.3xHA (y1w*;;PBac{UAS-CG17739-3xHA}VK00033/ TM3,Sb1Ser1)*. Gateway LR clonase II (ThermoFisher # 11791020) was used to make pGW-RELN or pGW-CG17739 from the pGW-attB-3xHA destination vector [91] and pENTR223 or pDONR223 vectors containing the cDNA sequences. pENTR223.1-RELN cDNA was commercially available, clone 100068963 (Accession Number: BC172269.1). pDONR223-CG17739[ter] was derived from pOT2-CG17739 cDNA, clone GH02025 (DGRC Stock 7692; https://dgrc.bio.indiana.edu//stock/7692; RRID:DGRC_7692; GenBank: NM_136913.3). The *CG17739* sequence containing the native stop codon was PCR amplified using Q5 Hot Start High-Fidelity 2X Master Mix (New England Biolabs # M0494) and primers which added flanking attB sites (forward primer 5’-GGGGACAAGTTTGTACAAAAAAGCAGGCTTCACCATGAAAGCTCTGGAAGGTTTCACTA-3’ and reverse primer 5’-GGGGACCACTTTGTACAAGAAAGCTGGGTCTCACCACCGCTCTGGATCATCGTTG-3’).

Gateway BP Clonase II (ThermoFisher # 11789020) was then used to insert the PCR product containing the CG17739 sequence into empty pDONR223. The stop codon was then removed using the Q5 Site-Directed Mutagenesis Kit (New England Biolabs # E0554; forward primer 5’-AGAGCGGTGGctgGACCCAGCTT-3’ and reverse primer 5’-GGATCATCGTTGCTGGTC-3’ to create the final open pDONR223-CG17739 vector. All clones were PCR and sequence confirmed. *CRIMIC* lines are generated as described in [35]. Briefly, the sgRNAs to target the genes and homology arms corresponding to 200 bps upstream and downstream of cut site (s) are synthesized in pUC57_Kan_gw_OK2 plasmid by Genewiz/Azenta. *attp-FRT-Splice acceptor-T2AGAL4-polyA-3XP3EGFP-polyA-FRT-attP* sequences of the corresponding phase are subcloned in the homology donor intermediate vector from pM37 vector (DGRC # 1517, 1519, 1520). sgRNAs used for *CG17739* are TATATACTTTAATCCTGTAGCGG and AACTATGTAATCATTACCAATGG.

### CG17739 Sequence analysis

The DIOPT-Ortholog prediction tool [26] and MARRVEL [92] were used to identify the most similar protein to human Reelin (*RELN*) in *D. melanogaster.* Protein BLAST tool [93] was also used to identify proteins with high similarities for human Reelin domains. Reelin aminoacid sequence and domains were obtained from UniProt (uniprot.org, P78509_RELN_HUMAN). Once the gene CG17739 from *D. melanogaster* (flybase.org, FBgn0033710) was identified, DIOPT-Ortholog prediction tool and MARRVEL were used to identify the most similar protein to CG17739 in *H. sapiens*. UniProt website was used to identify the protein domains of CG17739 (Q7K3Y9_DROME), F-spondin (Q9HCB6_SPON1_HUMAN) and Reelin (P78509_RELN_HUMAN) used to make the cartoon in Fig. 1A

### Protein Structure Prediction

The aminoacidic sequence of *Drosophila melanogaster* CG17739 gene was submitted to the Robetta protein structure prediction server. The RoseTTAFold method was used to generate the 3-dimensional structure of the protein [30]. The model was then processed with the Visual Molecular Dynamics software (VMD) to identify secondary structures and conformations of CG17739 protein.

### Myc-Drospondin construct

The aminoacidic sequence of CG17739-RA (FBtr0087893) was used for the design of the Myc-tagged-Drospondin construct. By site-directed mutagenesis the V5 tag of the pMT-Bip V5-His A vector was replaced by a Myc sequence (EQKLISEEDL) using the primers: Fw: 5’– AGCGAAGAGGATCTGACGCGTACCGGTCATCAT–3 and 5’– AATCAGTTTCTGTTCGAATTCCACCACACTGGACTAGTAGGTACC–3’ and the PFU ultra (Agilent). The cDNA for Drospondin without the signal peptide sequence and the stop codon was amplified using the clone GH02025 from *Drosophila* Genome Resource Center (DGRC). The forward primer 5’–GCGCGGAATTCATGCATTAGGGTCCCACCGG–3’ which contains EcoRI cut sites at the 5’ and the reverse primer: 5’–TTTTAGGCGCGCCCACCGCTCTGGATCATCG– 3’ which contains cut sites for BglII at the 5’ where used to amplify the cDNA of Drospondin.

### Drospondin stable transfection and production

S2 cells were maintained at 25°C in Schneider insect medium supplemented with 7% fetal bovine serum (FBS, Biological Industries, 04-127-1A) and 1% Penicillin/Streptomycin (P/S, GIBCO 15240-062) until the transfection. The Myc-Drospondin construct was transfected along with the plasmid pCoHygro in S2 cells using the calcium phosphate method following the instructions of Thermo Fisher Scientific Manual 0000656 recB0 catR69000. In parallel, S2 cells were transfected with the empty plasmid plus the plasmid pCoHygro. Two days after transfection the cell medium was replaced by selection media: Schneider insect medium supplemented with 7% FBS, 1% P/S and 300 μg/ml Hygromycin B (1,068,701, Invitrogen), until they arrest their division. Then, cells were diluted and seeded in a 96 well plate. After three days 48 cell colonies were selected and transferred to 24 well plates. By immunofluorescence (anti Myc antibody RRID: AB_325961, 1:100) (Figure S1A), using the same method described for brain immunofluorescence below, the best colony was selected and expanded. When S2 transfected cells reached 80% confluence the medium was changed to expression induction media: Schneider insect medium with 500µM CuSO_4_. 48 h hours later, the medium was collected for the first time and kept at 4°C, replaced by the same medium, and collected 24 h later again. This procedure was repeated one more time. The medium was diluted 5 times in PBS 1X and concentrated until 1/60 of the original volume was reached, using Amicon ultra-15 50kDa filters (Millipore). Drospondin concentration in the medium was estimated semi-quantitatively: BSA dilution curves and an aliquot of Drospondin-containing medium (and mock medium as control) were run through an SDS-PAGE. The gel was stained with Coomassie blue to visualize the Drospondin bands. The bands corresponding to Drospondin-encoded protein and BSA were digitalized to analyze their intensities in Fiji [140] (Figure S1B). The presence of Myc-Drospondin in the supernatant was corroborated by Western Blot using Anti-Myc 9E10 antibody (RRID: AB_2857941), 1:500 dilution.

### Evaluation of N-glycosylation Tunicamycin treatment

3.5 million of S2 cells stably transfected with Myc-Drospondin and the Mock cell line were plated in a 60 mm plate pretreated with Poly-D-lysine. The cells were kept at 25°C for 24 h in Schneider insect medium supplemented with 7% FBS, 1% P/S and 300 μg/ml Hygromycin B. Later, the medium was replaced by Schneider insect medium with 500µM CuSO_4_ and 5µg/ml Tunicamycin or the equivalent volume of DMSO in a final volume of 2ml. 48 h later, the supernatant was collected, centrifuged, diluted in PBS1X five times their volume and concentrated until 110 µL were obtained. All the recovered cells were lysed in 200 µL of Lysis Buffer (50 mM Tris, 50 mM NaCl, 20 mM EDTA and 1% Nonidet P-40) with protease inhibitors, centrifugated at 14000 rpm for 10 min at 4°C. 10 µg of the supernatant proteins and 45 µg of the cell lysates were charged in the electrophoresis acrylamide gel. The Western Blot was performed using Anti-Myc 9E10 antibody (RRID: AB_2857941) at 1:500 dilution and anti-beta actin antibody, 1:6000 (RRID: AB_449644), both diluted in blocking solution (5% milk in 0.1% Tween 20 PBS 1X).

### mRNA levels

Fifty adult male heads (0-3 days old) were used to extract the RNA and synthesize cDNA as previously described (Rojo-Cortés et al., 2022). The primers for qPCR reaction were designed to recognize two consecutive exons present in all the isoforms of the genes evaluated: Drospondin: 5’–ACCCTGCACATCAATCGAAGACG–3’ and 5’–GACCGCACTTTGTGGTG CATTC–3’, annealing at 60°C; LpR1: 5’–TGCACGAATGGAGCCTGCAT–3’ and 5’– GTGATGCGATCCTTGCACTGGT–3’, annealing at 60°C; LpR2: 5’– CTATGTCGTATACCGACGCTGC–3’ and 5’– CTGCGGCGTAAATGTGTGG–3’ annealing at 57.2°C. GAPDH was used as a reference gene: 5’ – CGTTCATGCCACCACCGCTA–3’ and 5’– CCACGTCCATCACGCCACAA–3’ annealing at 60°C. The qPCR reactions were performed using the Kit 5x HOTFIREPol EvaGreen qPCR Mix Plus (Solis BioDyne 08-24-00001) in 96 wells MicroAmp Optical reaction plate in Quant Studio3 Applied Biosystems (Thermo Scientific) apparatus. Primer efficiency was calculated from serial dilution as 10^-1/slope^. The relative expression was calculated as [primer efficiency for GAPDH^CT^ ^average^ ^among^ ^replica^]/ [primer efficiency for the gene of interest^CT^ ^average^ ^among^ ^replica^].

### Survival assay

Three groups of twenty (ten females and ten males) 0-2 days old flies per genotype were placed in fresh tubes with standard food. Flies were changed to a vial containing fresh food every 2-3 days. The number of dead flies was registered daily.

### Fly brain immunofluorescence

Larval L3, Pupae P9-10 and 0-3 days old adult male flies were dissected in PBS1X and fixed in Paraformaldehyde solution 4% in 0.3% PBS-Triton (PBS-T) for 20 min. Then, the brains were rinsed with PBS-T. The tissue was blocked (solution: 3% Normal Goat Serum, NGS; 0.03% Sodium Azide; in PBS-T) for 1 h. Later, the brains were incubated overnight with primary antibodies at 4°C under agitation. After rinses with PBS-T, a 3 h incubation with secondary antibodies was carried out at room temperature (RT). Brains were then washed and mounted with ProLong Gold (Invitrogen). The primary antibodies used were 1D4-FasII antibody (RRID: AB_528235) 1:20; 8D12-Repo antibody (RRID: AB_528448) 1:20; Elav-7E8A10 antibody (RRID: AB_528218) 1:50 and HA antibody (RRID: AB_1549585). The secondary antibodies were Alexa Fluor 568 goat anti-mouse (RRID: AB_144696) 1:500; Alexa Fluor 647 Goat anti-mouse (RRID: AB_2633277), 1:500; Alexa Fluor 555 Goat anti Rabbit (RRID: AB_2535849), 1:500; Alexa Fluor Goat anti Rat 647 (RRID: AB_141778) 1:500. The images were acquired using a Z-step size of 1.5 μm until the whole brain was covered. MB morphology was evaluated in both hemispheres from a Z project containing the entire MB area, in Fiji[94].

### Brain area

Z projection of each brain immunostained with anti-FasII was obtained using the maximum intensity option in Fiji. The brightest/contrast was adjusted to observe the border of the brains. Then, the perimeter of the central brain area was marked and measured three times per brain, using the freehand tool. The mean value of the three measurements was recorded as the area of a given brain. Brains were classified as “small” when their central brain area was smaller than the smallest area of the central brain recorded in the control strains.

### Primary neuronal culture from pupae

Neuronal cultures from pupae were performed as previously reported [17, 95]. Briefly, 9-10 pupal brains were dissected in dissection solution (DS: 6.85 mM NaCl; 0.27 mM KCl; 0.009 mM Na_2_HPO_4_; 0.001 mM KH2PO4; 0.2772 mM HEPES; pH 7.4). The brains were treated for 30 min with 9.6 U/ml of Papain (Worthington, LS 03126) at RT and then disgregated mechanically. The cell suspension obtained was placed over Laminin-Concavalin A coated coverslips in presence of DMEM/F12 culture medium (Gibco, 12400-016) supplemented with 100μg/ml Apo-Transferrin; 30 nM Selenium; 50 μg/ml Insulin; 2.1 mM 20-Hydroxyecdysone; 20 ng/ml Progesterone; 100 μM Putrescine (all supplements from Sigma-Aldrich); and 1% Antibiotic/Antimycotic (Gibco 15240062). The next day, conditioned media obtained from astrocytes cultured in Neurobasal medium supplemented with B27 (CNBM/27), was added to the brain culture in 1:3 ratio.

### Treatments of primary brain cultures for Sholl Analysis

Primary neuronal cultures from pupal brains were treated with different concentrations of Drospondin or Reelin, as in [17]. 48 h later the cultures were fixed with PFA 4%, sucrose 4% in PBS1x, for 20 min. Then, cell cultures were incubated for 15 min with 0.15M Glycine and washed three times with PBS-T and one time with distilled water. The coverslips were mounted using Fluoromount-G. Microphotographs were acquired using Nikon Ti 60x epifluorescent microscope. Sholl analyses were performed using the plugin SNT in Fiji [96].

### Sleep patterns and locomotor activity

Locomotor activity was measured using the *Drosophila* Activity Monitoring System (DAM, TriKinetics Inc, USA). 4-6 days old male flies were collected, separated by phenotype and transferred to glass capillaries containing 3% sucrose in a 1.5% agar prepared in distilled water. Flies were placed in an incubator at 25°C and 75% humidity, connected to the monitoring system (TriKinetics, Waltham, MA, United States) under 12 h:12 h light-dark cycles for 7 days.

The sleep analysis was performed in R using the Rethomics framework [97] after six days of baseline recordings. Sleep is defined as a continuous period of inactivity lasting 5 min or more [98, 99]. The activity counts to actograms profiles were calculated as the number of beam crossings each 60 min (Fig 7).

### Social distance essay

Procedure was as in [58, 100]. Briefly, groups of 30 to 40 male flies (3-5 days old) were placed in the triangle arena, at 50% humidity, 25°C. The arena was placed vertically, and immediately after all flies reached the top of the triangle, the arena was shifted to a horizontal position. After 45 min, three consecutive pictures were taken of the arena (<1 seg between them), to record the position of each fly. The plug-in “nearest neighbor” [100] was used to calculate the distance between each fly and their nearest neighbor. The average value of distance to the nearest neighbor from each of the flies in one picture was averaged with the data from the other two pictures. This number represents one value in the graph.

### Individual locomotor activity in a circular arena

As previously reported [58, 101]. Briefly, flies were aged to the corresponding age window. 20 h before the start of a given experiment, the males of the desired genotype were collected using cold anesthesia. On the day of the experiment, groups of 4-6 flies were placed over a white acrylic and covered with a 39 mm petri dish; all the procedure was carried out under cold anesthesia. Then, the flies were moved to the behavior room (22-24°C, 40-60% humidity) and were left there for 40 min (acclimation period). Later, single flies were transferred to a new white acrylic covered with a 35mm diameter petri dish and placed over a thermic block at 25°C. The arena was surrounded by a 10 cm high white wall. The basal locomotor activity was recorded for 5 min using a digital camera placed 60 cm over the arena. The software PylonViewer was used to record the videos, and IowaFLI Tracker [102] was used to analyze them.

### Data analysis

The software GraphPad Prism was used for statistical analyses, except for the survival assay which was analyzed using R 2022.07.1 and the packages “survival” and “survminer”.

## ACKNOWLEDGMENTS

We thank Juan Bonifacino and Evelyn C. Avilés for comments and suggestions on earlier versions of this manuscript. We would like to acknowledge the Bloomington Drosophila Stock Center for fly lines and the help provided by the Advanced Microscopy Facility UC and the sequencing unit UC: CONICYT – FONDEQUIP EQM150077. This work was supported by FONDECYT grants 1200393 (to MPM), 1191424 and 1231685 (to CO), and 1141233 and 1231556 (to JMC). FR-C, CBR, PA-H and IA-T were supported by ANID Doctoral fellowships N° 21180582, 21240933, 21221773 and 21240414, respectively. LDG was supported by the Postdoctoral Fellowship in Alzheimer’s Disease Research from the BrightFocus Foundation. OK was supported by NIA Grant R24OD031447. OK was supported by NIH ORIP grant R24OD031447. This work utilized reagents made available by the Drosophila Genomics Resource Center (NIH Grant 2P40OD010949).

## AUTHOR CONTRIBUTIONS

Conceptualization, MPM, JMC and FR-C; Methodology, MPM, JMC, and FR-C; Formal Analysis, FR-C, CBR, NF-U, PA-H and IA-T; Investigation, FR-C, CBR, PA-H, IA-T and M-C.G-R; Resources, JMC, MPM, SB, LDG, OK and CO; Writing – Original Draft FR-C; Writing – Review & Editing, all authors; Funding Acquisition MPM, JMC and CO

